# Connecting Cryo-EM and Crystallographic Views of RNA Folding through Ionic Conditions and Structural Flexibility

**DOI:** 10.64898/2026.05.02.722415

**Authors:** Avijit Mainan, Serdal Kirmizialtin, Susmita Roy

## Abstract

Discrepancies between biomolecular structures resolved by cryo-electron microscopy (cryo-EM) and X-ray crystallography (XRD) often arise from differences in ionic conditions and construct design, yet their mechanistic impact on RNA folding remains unclear. In the SARS-CoV-2 frameshifting stimulatory element, cryo-EM and XRD structures reveal distinct pseudoknot conformations—a bent and a coaxially stacked state—complicating its structure–function relationship. Here, combining all-atom explicit-solvent simulation results with a structure-based electrostatic model, we show that Mg²⁺ ions drive transitions between these states by stabilizing long-range tertiary interactions and modulating local dynamical coupling involving the slippery site and stem 3. Energy landscape analysis reveals distinct folding pathways, while deletion of the slippery segment in crystallographic constructs alters intermediates and produces pathways inconsistent with single-molecule optical tweezer experiments. This study demonstrates how condition-dependent experiments encode complementary interaction-level information and how physics-based computational approaches integrate these to yield a coherent, mechanistic picture of RNA folding.

**TOC GRAPHICS:** 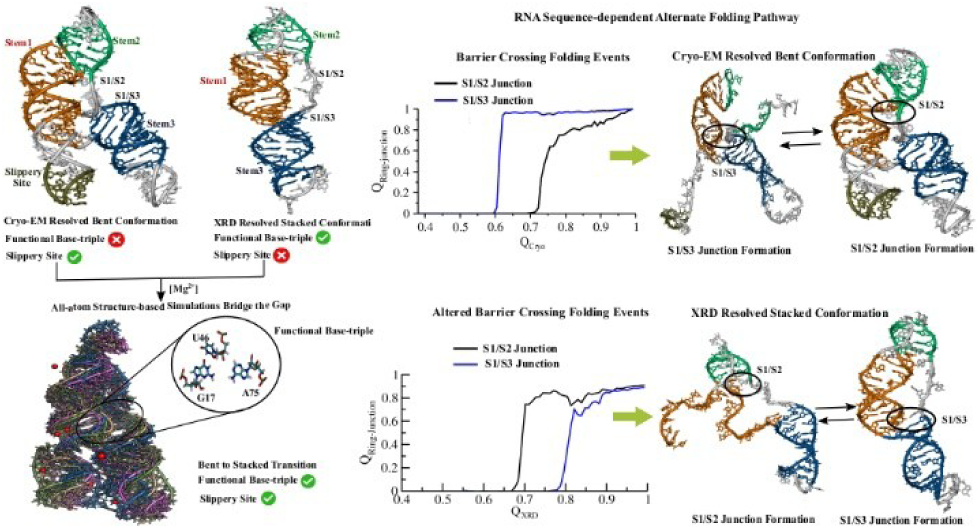

---

High-resolution structural determination of biomolecules has been revolutionized by techniques such as X-ray crystallography (XRD) and cryo-electron microscopy (cryo-EM). Despite their transformative impact, a growing body of work has highlighted systematic discrepancies between structures obtained by these methods, raising fundamental questions about how experimental conditions shape the observed conformational landscapes. XRD typically requires crystallization under non-physiological conditions, often involving high salt concentrations, precipitants, and lattice constraints that can restrict molecular flexibility or stabilize specific conformations^1,2^. In contrast, cryo-EM captures biomolecules in vitrified solution under near-native conditions and can resolve multiple conformational states within heterogeneous ensembles^3,4^. These differences frequently lead to variations in domain organization, tertiary contacts, and conformational heterogeneity across a wide range of systems, including globular proteins, membrane proteins, intrinsically disordered proteins (IDPs), and nucleic acids.

Such discrepancies are particularly pronounced in RNA, where folding is governed by a delicate interplay between electrostatic interactions, sequence context, and ionic conditions. RNA pseudoknots (PKs), in particular, are inherently dynamic and capable of adopting multiple conformational states, with their stability and folding pathways strongly modulated by divalent cations such as Mg²⁺ ion^5–7^. However, resolving these conformational ensembles experimentally remains challenging. Early studies combining graph-theory-based modelling with SHAPE experiments revealed that the SARS-CoV-2 RNA PK adopts multiple conformations that influence frameshifting efficiency^8,9^. More recent work, integrating atomistic simulations with experimental data, further demonstrated that chemical modifications can significantly affect RNA structural geometries, emphasizing their impact on interpreting experimental structures^10,11^.

Structural studies of ribosomal RNA, viral RNA elements, and riboswitches have repeatedly revealed inconsistencies between XRD and cryo-EM structures. Early comparisons showed that crystal packing forces and construct truncations can bias RNA toward compact or coaxially stacked conformations, whereas cryo-EM structures—often determined in more native-like environments—capture bent or alternative states that reflect functional dynamics^12,13^. Complementary solution-based techniques, such as single-molecule FRET and optical tweezers, further highlight folding intermediates and pathways that are frequently absent in crystallographic models^14,15^.

Importantly, these discrepancies are not solely methodological but are also influenced by sequence modifications and environmental conditions. Crystallographic constructs often involve truncations of flexible regions or sequence alterations to facilitate lattice formation, while differences in ionic conditions—particularly Mg²⁺ concentration—can significantly alter tertiary interactions and long-range coupling within RNA. Although such effects have long been recognized, there remains a lack of explicit demonstration and mechanistic understanding of how condition-dependent variations may modulate RNA structure, dynamics, and ultimately function.

The frameshifting stimulatory element (FSE) of SARS-CoV-2 provides a compelling system to address this gap. Recent high-resolution structures obtained by cryo-EM and XRD reveal distinct conformational states of its pseudoknot (PK), including bent and coaxially stacked arrangements, respectively. One conformation was determined by cryo-electron microscopy (cryo-EM)^16^, while the other was resolved by X-ray crystallography (XRD)^17^, with the crystallographic structure obtained in complex with a chaperone. These structures are further supported by the ribosome-bound structure of the PK ^18^ and by solution small-angle X-ray scattering (SAXS) studies of the PK in monovalent salt conditions ^19^. All structures share a common global architecture comprising three stems (stem1, stem2, and stem3) connected by two ring junctions (S1/S2 and S1/S3), through which the 5′ end of the RNA is threaded, forming the characteristic FSE topology (**Figure 1**).

**Figure 1.**
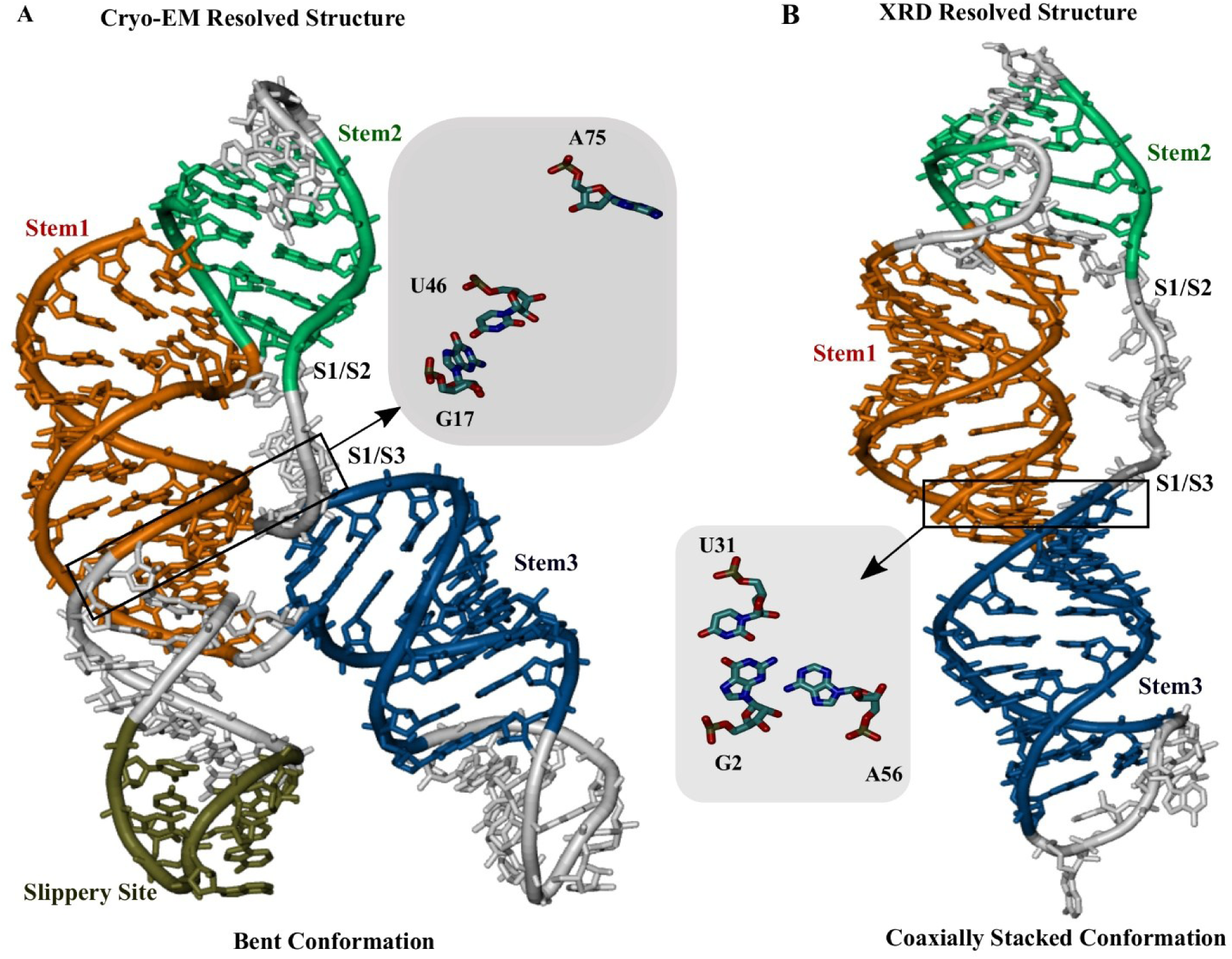
Tertiary structures of the SARS-CoV-2 mRNA frameshifting element resolved by cryo-electron microscopy (cryo-EM) and X-ray crystallography (XRD). (A) Bent conformation of the three-stemmed SARS-CoV-2 pseudoknot resolved by cryo-EM, which retains the slippery site (PDB ID: 6XRZ). At the S1/S3 junction, residues G17 and U46 form a base pair, while A75 does not participate in base-triple formation. (B) Coaxially stacked conformation of the three-stemmed SARS-CoV-2 pseudoknot resolved by XRD (PDB ID: 7MLX). At the S1/S3 junction, residues G2, U31, and A56 form a stable base-triple interaction.

In the crystal structure, the three stems adopt a vertically stacked configuration stabilized by a G2–U31–A56 base-triple interaction at the S1/S3 junction (**Figure 1B**), a structural feature proposed to be critical for programmed ribosomal frameshifting^17^. In contrast, this base triple is not observed in the cryo-EM models, although its absence may reflect limitations in resolution rather than a genuine lack of tertiary interactions. The G2-U31-A56 base triple at the S1/S3 junction is expected to stabilize the vertical conformation further and cannot be fully accommodated in the bent geometry resolved by cryo-EM. Specifically, in the cryo-EM structure, G17 and U46 are positioned close enough to form a base pair, whereas A75 remains more than 10 Å away (**Figure 1A**). The residue numbering of this base triple differs between the two structures due to the absence of the slippery site in the crystallographic construct. Nevertheless, the G2–U31–A56 base triple observed in the XRD structure corresponds to the same nucleotides as G17–U46 and A75 in the cryo-EM structure.

These differences raise critical questions regarding the structural determinants of functional RNA states and the extent to which experimental conditions—such as ionic environment and construct design—control the observed conformations. In particular, the omission of flexible elements, such as the slippery site in crystallographic constructs, may alter folding pathways and stabilize non-native intermediates. These complementary limitations highlight the need for integrative approaches. In this context, all-atom molecular dynamics (MD) simulations, particularly when combined with a physics-based electrostatic model, provide a powerful framework to bridge experimental observations and reveal the underlying ion-dependent and sequence -dependent conformational dynamics of RNA.

Initially, to understand the salt concentration dependence of RNA structure, we perform classical explicit-solvent molecular dynamics simulations. The details of the explicit solvent simulation method have been described in the Supporting Information. We observe a Mg²⁺-dependent transition from the bent to the coaxially stacked conformation starting from the cryo-EM resolved structure as an initial structure, preparing a physiological ionic concentration (∼2 mM Mg²⁺; **Figure 2A, B**). To quantify this transition, we monitor the centre-of-mass distance between the slippery site (residues 4–10) and the stem3 bulge region (residues 52–67) at ∼100 mM K⁺, both in the absence and presence of Mg²⁺. The broad probability distribution of the centre-of-mass distance between the slippery site and stem3 in Figure 2A reflects a heterogeneous ensemble of structures (**Figure S1**). Upon addition of 2 mM Mg²⁺, the initially bent conformation undergoes a clear transition to a coaxially stacked state. Interestingly, this transition is accompanied by a closer approach of a base-triple interaction (G17-U46-A75) as shown in **Figure S2,** which was not resolved in the cryo-EM experiment.

**Figure 2.**
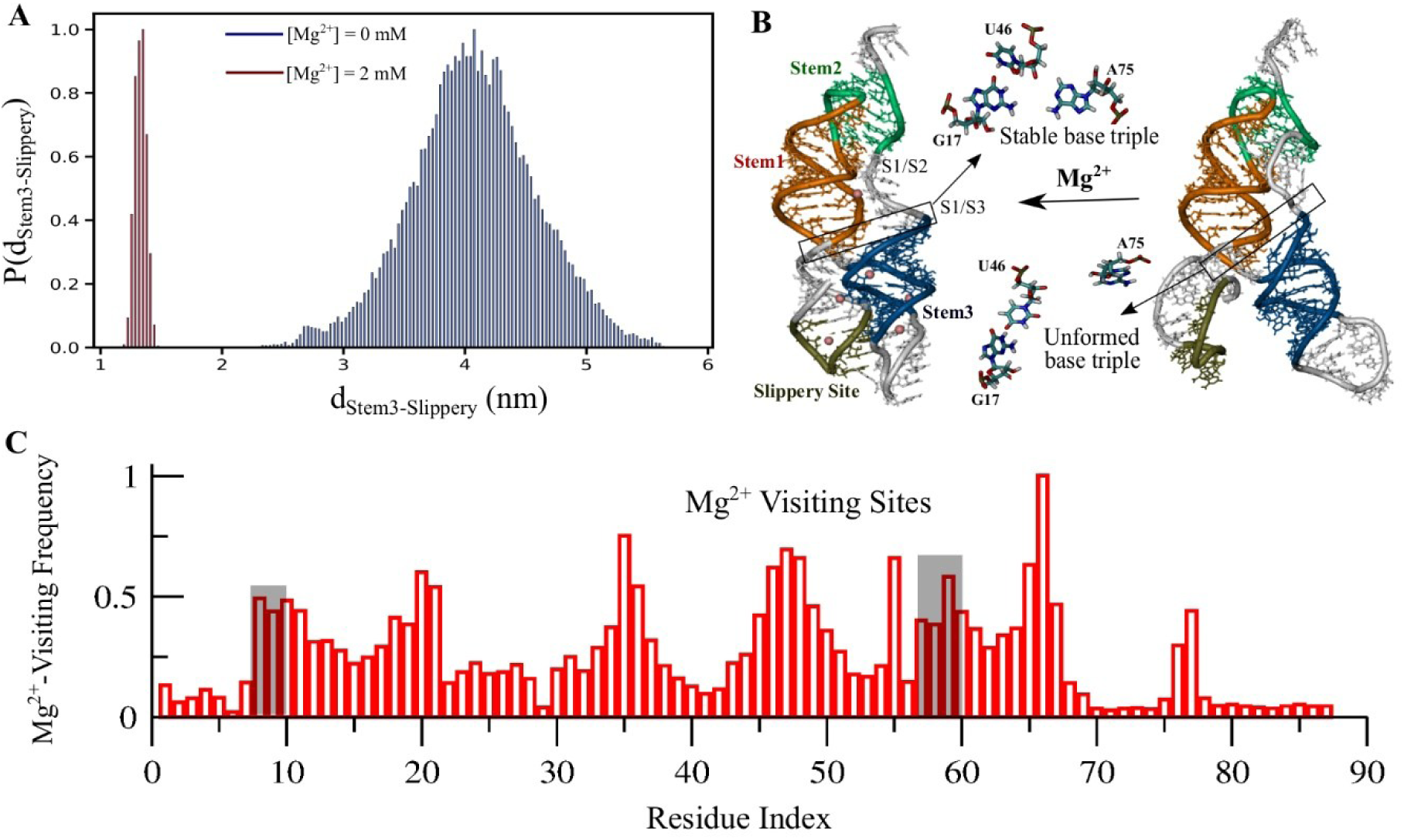
All-atom MD simulations results of the RNA in the absence and presence of Mg^2+^ ions. (A) Distance distribution between stem3 and the slippery site at [Mg^2+^] = 0 mM and [Mg^2+^] = 2 mM conditions. (B) Highlighting Mg²⁺-dependent structural rearrangements, with representative snapshots illustrating the transition from a bent conformation lacking a stable base triple to a coaxially stacked conformation, facilitates a closer approach to a base-triple interaction (G17-U46-A75). (C) Site-visiting frequency of Mg²⁺ ions around phosphate groups along the RNA sequence.

To elucidate the specific role of Mg²⁺ ions in driving this conformational transition, we quantify the site-visiting frequency of Mg²⁺ ions around each phosphate group of the RNA. While multiple regions along the RNA display highly frequent Mg²⁺ visiting sites, two dominant hotspots emerge near the slippery site and stem3 (**Figure 2C**). Complementary to the site-visiting analysis, we searched for Mg binding motifs from the cryo-EM structure using a recently developed stereochemistry-guided approach, CatWiz ^20^. Using CatWiz, we identify a well-defined inner-sphere Mg²⁺ binding site near the slippery site (**Figure S3**), which overlaps with simulation results (**Figure 2C**). A representative structural snapshot highlighting this binding site is shown in **Figure S3B**. It is important to note that CatWiz classifies binding when it matches the set geometric and distance criteria described only for direct/inner-shell coordination ^20^. In MD simulation visiting frequency analysis, on the other hand, we consider Mg^2+^ ions within the cut-off distance from the phosphate backbone, accounting for both inner and outer-sphere (solvent-separated) binding/coordination modes. We adopt the criteria based on our early studies^21–23^, where we found the critical role of outer-sphere (solvent-separated, hexa-hydrated Mg^2+^) linking the long-range tertiary connection in an RNA fold.

To elucidate structural details, we also compute contact probability maps (**Figure 3A**). In these maps, the upper triangular region corresponds to the Mg²⁺-free condition, whereas the lower triangular region represents the 2 mM Mg²⁺ condition. Contacts were calculated at the all-atom level and represented in a residue-wise manner. The resulting maps reveal that, in the absence of Mg²⁺, interactions between the slippery site and stem3 are not present, while at 2 mM Mg²⁺, these interactions become significantly enhanced. Together, these results demonstrate that the bent-to-stacked conformational transition is driven by Mg²⁺-dependent stabilization of tertiary interactions. These observations also suggest that the S1/S3 junction plays a central role in governing the dynamic interconversion between bent and coaxially stacked conformations, with potential consequences for frameshifting efficiency.

**Figure 3:**
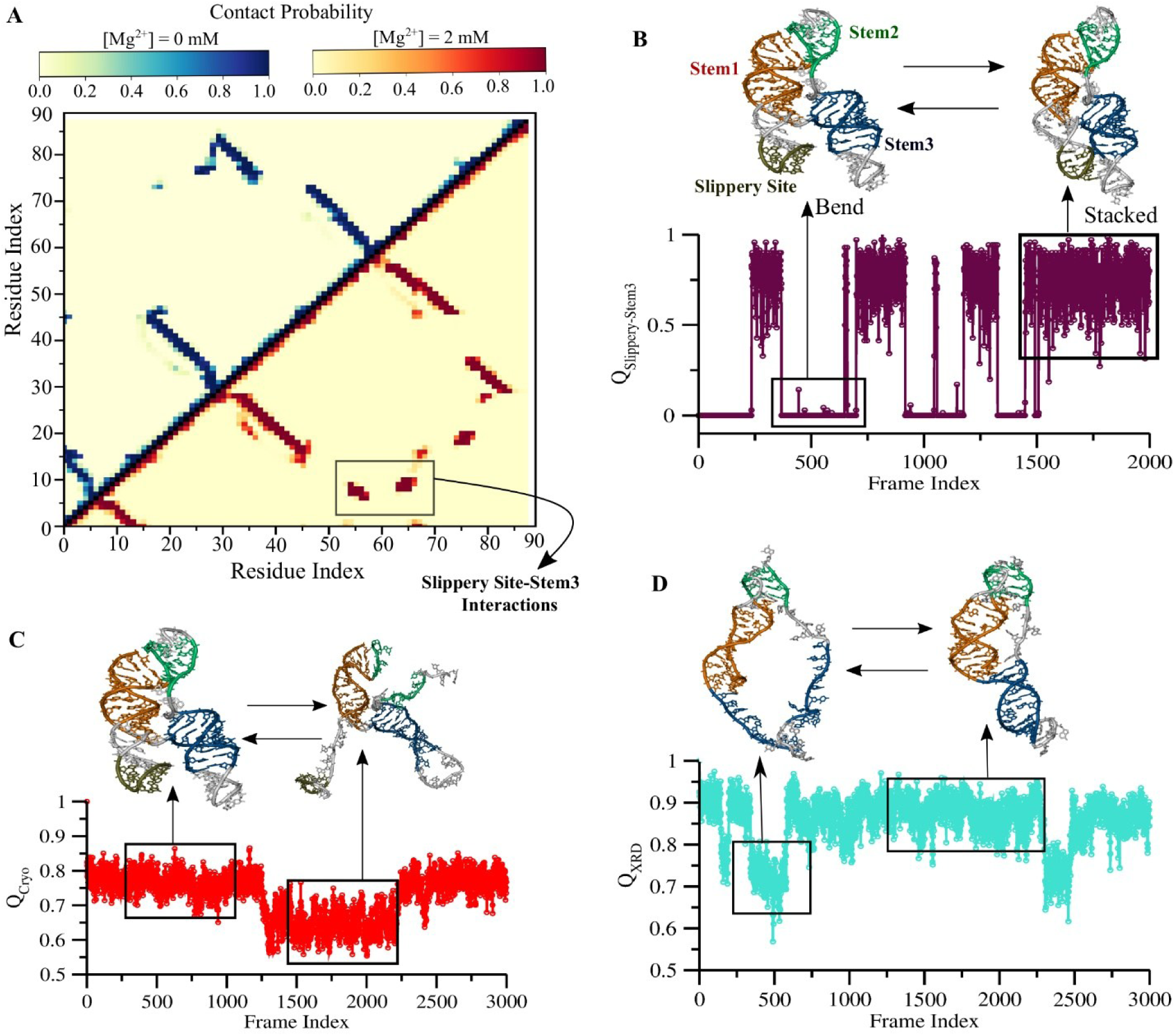
Explicit-solvent–guided STEM framework capturing major conformational breathing between the cryo-EM and XRD structures of the SARS-CoV-2 RNA. (A) Residue-wise contact probability maps from all-atom RNA simulations under Mg²⁺-free and 2 mM Mg²⁺ conditions; the upper triangle corresponds to the Mg²⁺-free condition, while the lower triangle corresponds to 2 mM Mg²⁺. (B) STEM simulations illustrating the transition from the bent to the coaxially stacked conformation, along with representative structural snapshots. (C) Time evolution of the fraction of native contacts relative to the cryo-EM structure (Q_Cryo_), capturing the transition from a stem2-open to a stem2-closed state. (D) Time evolution of the fraction of native contacts relative to the crystal structure (Q_XRD_), capturing the transition from a stem3-open to a stem3-closed state.

Explicit-solvent molecular dynamics simulations suggest that the transition from the bent to the coaxially stacked conformation is modulated by Mg²⁺ concentration. However, the accessible timescales of all-atom simulations are insufficient to capture reversible bent-to-stacked transitions and fully explore the folding landscape of the RNA. To overcome the limitation of simulating long-time scale associated phenomena, our group has recently developed a Structural Topology-based Electrostatic Model (STEM) integrating salt-associated dynamic counter-ion condensation phenomena and explicit ion-ion correlation involving Mg^2+^ mediated interaction^21,22,24^. To implement counter-ion condensation phenomena into an all-atom structure-based RNA simulation method, Generalized Manning Counter-ion Condensation (GMCC) theory^25^ is used in STEM. GMCC theory extends the classical Manning condensation theory^26,27^ by considering counterion condensation for polyelectrolyte systems of irregular shape and charge distribution. STEM addresses the physiological salt environment under which RNA is exposed to a mixture of monovalent (concentration range ∼100-500 mM) and divalent salts (∼1-2 mM) ^28,29^. Explicit treatment of Mg^2+^ is crucial for directly controlling RNA tertiary packing, accounting for spatial ion-ion correction, whereas intermediate KCl concentration is treated implicitly for computational efficiency using the GMCC model. Because of the implicit treatment of salt buffer, this structural Topology-based Electrostatic Model (STEM) is computationally economical and can efficiently sample a large phase space of a dynamic system like RNA along with its dynamic ion environment. The details of this approach are provided in the Supporting Information and our early works^22^.

While the current form of STEM potential effectively apprehends the dynamic cooperativity of ion condensation and maintains the site specificity between implicit K^+^ and explicit Mg^2+^ ions ^30^, the explicit Mg^2+^ are primarily an outer-sphere of type (accounting for the hydrodynamic radius of hexahydrated Mg^2+^). We are working to improve STEM potential to incorporate dynamic ion-exchange of inner-outer-sphere coordination mode, accounting for both inner and outer-sphere coordination of Mg^2+^. In this work, the inner-sphere effect has been implicitly included into the contact-potential of the formal form of STEM potential, in terms of inner-sphere Mg^2+^ mediated tertiary contact information. This non-native all-atom-level microscopic contact information we obtain from equilibrated explicit solvent simulation analyses (**Figure 3A**).

From explicit solvent simulations, we observe that the closure between the slippery site and stem3 involves long-range interactions mediated by Mg²⁺ ions. We explicitly incorporated these interactions into the formal form of STEM Hamiltonian (Eq. 1and Eq. 2). To guide this incorporation, we analyzed contact probability maps obtained from explicit-solvent simulations under Mg²⁺-free and 2 mM Mg²⁺ conditions (**Figure 3A**). Contacts that emerge as unique contacts specifically in the presence of Mg²⁺ were identified as Mg²⁺-mediated unique interactions and subsequently included in the Hamiltonian. In particular, this procedure captures the formation of long-range contacts between the slippery site and stem3, which are essential for describing the Mg²⁺-dependent conformational transition. The methodology for calculating Mg²⁺-mediated contacts has been described in detail in our recent work^31^.

The total Hamiltonian employed in the STEM framework is expressed as

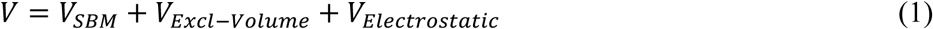

where 𝑉_𝑆𝐵𝑀_ represents the all-atom structure-based model potential^32–34^, which is further decomposed into native and Mg²⁺-mediated interactions as

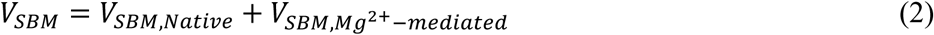

Here, 𝑉_𝐸𝑥𝑐𝑙−𝑉𝑜𝑙𝑢𝑚𝑒_ accounts for excluded-volume interactions involving explicitly treated Mg²⁺ ions, while 𝑉_𝐸𝑙𝑒𝑐𝑡𝑟𝑜𝑠𝑡𝑎𝑡𝑖𝑐_represents all electrostatic interactions within the nucleic acid system. Additional details of the STEM Hamiltonian are provided in the Supporting Information.

The Mg²⁺-dependent slippery site–stem3 interactions are therefore incorporated into the STEM Hamiltonian (𝑉_𝑆𝐵𝑀,𝑀𝑔_^2+^_−𝑚𝑒𝑑𝑖𝑎𝑡𝑒𝑑_) to enable the model to capture the observed conformational transition. Time evolution of the inter-segment fraction of native contacts between stem3 and the slippery site shown in **Figure 3B** reveals a clear transition from the bent to the coaxially stacked conformation (see also Supporting Information, **Movie S1**). In this manner, explicit-solvent MD simulations provide a quantitative basis for parameterizing and benchmarking the STEM framework to capture the Mg²⁺-driven bent-to-stacked transition.

With the STEM Hamiltonian trained for this RNA, we next compare the conformational dynamics of the cryo-EM and crystal structures. For the cryo-EM structure, equilibrium STEM simulations reveal frequent transitions corresponding to stem2–open (Q_Cryo_ ≈ 0.65) and stem2–closed states (Q_Cryo_ ≈ 0.80) (**Figure 3C**). In contrast, simulations initiated from the crystal structure exhibit transitions between stem3–open (Q_Crys_ ≈ 0.65) and stem3–closed states (Q_Crys_ ≈ 0.85) (**Figure 3D**). Interestingly, during the stem3 opening–closing transitions in the crystal structure, the stable S1/S3 base-triple interaction becomes distorted (**Figure S4** in Supporting Information). This destabilization is likely a consequence of truncating the slippery site in the crystallographic construct. Given that the S1/S3 base triple has been implicated as a key structural element in programmed ribosomal frameshifting^17^, destabilization of this interaction at physiological Mg^2+^ concentration range may influence frameshifting efficiency.

To investigate the folding mechanism of the SARS-CoV-2 FSE and its sensitivity to sequence construct (with and without slippery segment), we compute the folding free-energy landscape as a function of the global fraction of native contacts and compare the free energy mechanism that accounts for the bent cryo-EM–derived conformation (Q_Cryo_) and the coaxially stacked XRD-derived conformation (Q_Crys_). Free-energy profiles are obtained using umbrella sampling integrating with the STEM Hamiltonian, which enables the exhaustive exploration of the entire phase-space by partitioning the collective variable into overlapping windows and reconstructing the full free-energy landscape from the resulting histograms using the Weighted Histogram Analysis Method (WHAM). Details of the umbrella sampling protocol and free-energy reconstruction are provided in the Supporting Information.

For the cryo-EM structure, the folding free-energy profile exhibits three well-separated minima, corresponding to three distinct conformational ensembles: (i) a folded state characterized by a stem2–closed conformation (F), (ii) an intermediate ensemble in which stem2 is open (I), and (iii) an unfolded ensemble (U) (**Figure 4A**). As stem1–stem2 (S1/S2) and stem1–stem3 (S1/S3) junctions are key tertiary structural elements implicated in programmed ribosomal frameshifting, we next examine the hierarchical order of sequence formation along the folding free energy profile, especially to identify major barrier-crossing events. Analysis of the tertiary folding pathway, first accounting for the cryo-EM structure, reveals a hierarchical folding process in which the S1/S3 junction forms during the first barrier-crossing event (Q_Cryo_ ≈ 0.62), followed by formation of the S1/S2 junction during the second barrier-crossing event (Q_Cryo_ ≈ 0.75), as illustrated in **Figure 4B**.

**Figure 4.**
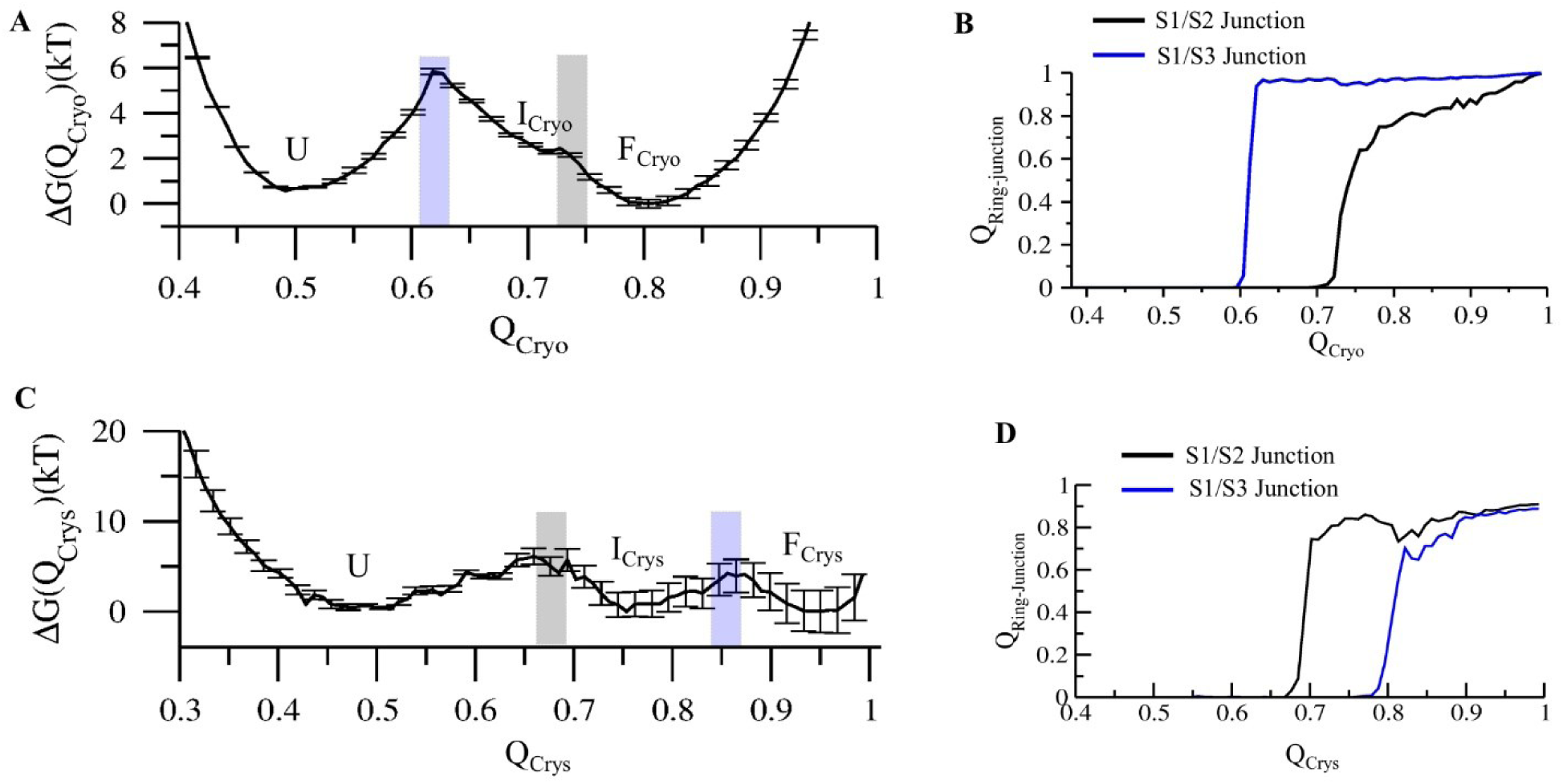
Folding free-energy landscapes of the SARS-CoV-2 pseudoknot evaluated from the STEM framework for the cryo-EM and crystal structures. (A) Folding free-energy profile of the cryo-EM structure, exhibiting three distinct minima corresponding to the unfolded state (U), an intermediate stem2–open state (I_Cryo_), and the fully folded stem2–closed state (F_Cryo_). (B) Sequential formation of the two junctions (S1/S2 and S1/S3 junction) as a function of the global native contact fraction (Q_Cryo_) for the cryo-EM–resolved structure. (C) Folding free-energy profile of the crystal structure, displaying three distinct minima corresponding to the unfolded state (U), an intermediate stem3–open state (I_Crys_), and the fully folded stem3–closed state (F_Crys_). (D) Sequential formation of junctions S1/S2 and S1/S3 as a function of Q_Crys_ for the XRD-resolved structure.

In contrast, the folding free-energy profile calculated accounting for the crystal structure also displays three barrier-separated minima, but with distinct structural assignments: (i) a folded ensemble corresponding to a stem3–closed conformation (F), (ii) an intermediate ensemble in which stem3 remains open (I), and (iii) an unfolded ensemble (U) (**Figure 4C**). Examination of the tertiary junction formation along this free-energy landscape reveals a different folding hierarchy. Specifically, the S1/S2 junction forms during the first barrier-crossing event (Q_Crys_ ≈ 0.70), followed by formation of the S1/S3 junction during the second barrier-crossing event (Q_Crys_ ≈ 0.85), as shown in **Figure 4D**. The nature of stability can be altered by varying temperatures. The temperature sensitivity of the free energy profile is shown in **Figure S5**.

Taken together, these results demonstrate that the cryo-EM and crystal structures are associated with distinct folding free-energy landscapes and intermediate states. The order of tertiary junction formation differs between the bent structure with a slippery segment and coaxially stacked conformations without a slippery segment, highlighting how structural context and topology reshape the folding pathways of the SARS-CoV-2 FSE.

The crystallographic study, that resolves a coaxially stacked conformation in the absence of the slippery site, compares its folding pathway with that obtained from optical tweezers experiments, while explicitly noting that pathway correspondence from their construct without slippery segment requires further validation^17,35^. Our free-energy landscape analysis and characterization of tertiary junction formation already indicate that the bent and coaxially stacked conformations follow distinct folding pathways^30^. Moving beyond 1D pathway, to further explore the possibility of multiple pathway mechanism associated with this RNA folding, we performed multiple temperature-dependent unfolding and refolding simulations of the SARS-CoV-2 frameshifting stimulatory element (FSE) by using the STEM framework, separately initiating from the bent (without slippery) and stacked (with slippery) structures. All resulting trajectories were concatenated to construct three-dimensional free-energy surfaces using the fractions of native contacts of stem1, stem2, and stem3 as collective variables.

To resolve the underlying kinetic pathways, we applied a Markov state model (MSM) framework to the unfolding–refolding trajectories. The methodological details are provided in the Method section (see Supporting Information). For the cryo-EM–resolved structure, the MSM analysis reveals a dominant single folding pathway that proceeds through a stem2–open intermediate state (**Figure 5A**). Especially, this pathway connecting the landmark intermediate, closely aligns with that of the folding pathway inferred from optical tweezers experiments^35^, consistent with the presence of the slippery site in both the cryo-EM construct. In contrast, analysis of the crystal structure reveals two alternative folding pathways that proceed through stem3 opening (**Figure 5B**). It is important to note that the folding pathway associated with the crystal structure exhibits greater heterogeneity, characterized by five dominant conformational state population. For each of these population cluster, one representative snapshot is shown along the characteristic pathway. The cryo-EM–derived folding pathway is comparatively less heterogeneous and is dominated by three state populations. Importantly, within the conformational ensemble sampled by the crystal structure, none of the populated intermediate states correspond to a stem2–open configuration, which constitutes a key intermediate observed in optical tweezers experiments and our present work.

**Figure 5.**
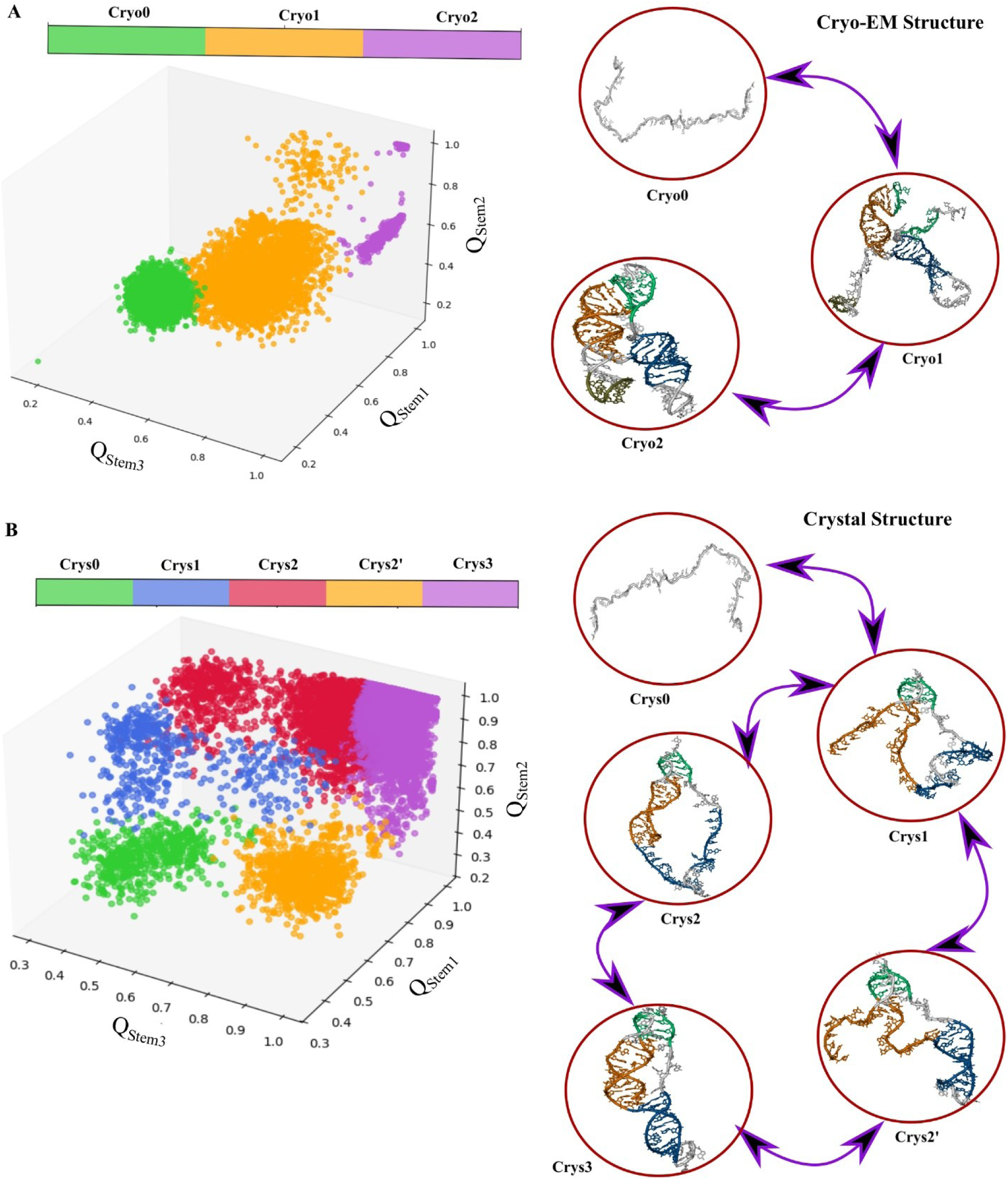
Three-dimensional free-energy surfaces (FES) and Markov state model–based clustering analysis of SARS-CoV-2 pseudoknot folding obtained from STEM kinetic simulations, comparing crystal and cryo-EM structures. (A) Three-dimensional FES constructed using the fraction of native contacts of stem1, stem2, and stem3 as order parameters, showing state populations and dominant folding pathways for the crystal structure. (B) Corresponding 3D FES and folding pathways for the cryo-EM–resolved structure, constructed using the same order parameters as in (A).

These findings demonstrate that, in the absence of the slippery site, the folding pathway of the coaxially stacked conformation diverges fundamentally from that observed experimentally. Consequently, truncation of the slippery site in the crystallographic construct alters the folding landscape and hampers direct comparison with experimentally measured folding pathways.

For SARS-CoV-2 FSE, each experimental approach offers distinct advantages and limitations: cryo-EM captures a structure that retains the functionally critical slippery site, which is essential for programmed ribosomal frameshifting but too flexible to be resolved in crystallographic constructs, whereas XRD provides high-resolution structural detail, revealing base-triple interactions at the S1/S2 and S1/S3 junctions that are not resolved in cryo-EM, likely due to resolution constraints^16,17^ (**Figure 1**). In this study, we sought to bridge the gap between these limitations coming from different experimental methods by using our computational approach.

To reconcile these, we combined explicit-solvent molecular dynamics simulations with our recently developed STEM framework. By systematically comparing the bent cryo-EM–derived conformation, which includes the slippery site, with the coaxially stacked crystal structure, which lacks this element, we demonstrate that these states are not merely alternative static snapshots but are embedded within distinct thermodynamics.

Explicit-solvent simulations reveal that the transition from the bent to the coaxially stacked conformation is driven by Mg²⁺ ions, which preferentially localize near the slippery site and stem3, promote Mg²⁺-mediated long-range interactions (**Figure 2**). These ion-mediated interactions stabilize key tertiary contacts, including the closer approach to a base-triple interaction (G17-U46-A75) as shown in **Figure 2B and Figure S2**, that is not resolved in the cryo-EM structure but emerges dynamically under physiological Mg²⁺ concentrations.

Using STEM simulation approach, we further dissect the free-energy profiles and folding pathways of the bent and stacked conformations. Although both landscapes feature unfolded, intermediate, and folded ensembles, the identities of the intermediates and the hierarchy of tertiary junction formation differ noticeably (**Figures 3, 4**). The bent cryo-EM structure folds through a stem2–open intermediate with early S1/S3 junction formation, followed by S1/S2 closure, whereas the crystal structure follows a pathway dominated by stem3 opening and closing with a reversed order of junction formation. Consistent with these thermodynamic differences, kinetic analysis based on temperature-dependent unfolding–refolding simulations and Markov state modelling shows that the cryo-EM structure follows a dominant folding pathway that closely aligns with optical tweezers experiments^35^, reflecting the presence of the slippery site (**Figure 5A**). In contrast, the crystal structure exhibits greater pathway heterogeneity and samples alternative routes that lack the experimentally observed stem2–open intermediate, demonstrating that truncation of the slippery site fundamentally alters the folding free-energy landscape and pathway (**Figure 5B**).

Taken together, our results establish that Mg²⁺-mediated ion interactions and sequence completeness—particularly the presence of the slippery site and its long-range ion-mediated interaction with stem-3—jointly determine the thermodynamic stability, folding hierarchy, and kinetic pathways of the SARS-CoV-2 FSE. By integrating complementary experimental insights with a unified computational framework, this work reconciles apparent discrepancies between cryo-EM and crystallographic structures and underscores the importance of considering ionic environment and construct design when interpreting RNA structures. More broadly, our study demonstrates how physics-based computational approaches can bridge the limitations of individual experimental methods, providing a comprehensive view of ion-driven conformational transitions that regulate RNA folding pathways central to RNA’s critical function.

## ASSOCIATED CONTENT

### Supporting Information

The following files are available free of charge.

Details of all-atom explicit-solvent simulations; structural topology-based electrostatic model (STEM), folding free-energy simulation methods, Markov state model construction, STEM simulation details, figure of representative conformational ensemble of RNA under Mg²⁺-free conditions, Mg²⁺-induced closer approach of the G17–U46–A75 base-triple interaction, stereochemistry-guided Mg²⁺ binding sites, analysis of breathing dynamics in the XRD-resolved SARS-CoV-2 RNA and its impact on base-triple stability, temperature-dependent free-energy profiles; and references (PDF).

Movie S1: Transition from bent to coaxially stacked conformation obtained from STEM simulation (MP4).

## Notes

The source code for STEM can be downloaded from https://github.com/Avijit-Source-Codes/STEM. All other data and codes used in the analysis are available from the corresponding author to any researcher for the purpose of reproducing or extending the analysis, subject to a material transfer agreement with IISER-Kolkata, India.

The authors declare no competing financial interests.

## ACKNOWLEDGEMENTS

This work was supported by the Department of Biotechnology (DBT) (Grant No. BT/12/IYBA/2019/12 and BT/PR40192/BTIS/137/69/2023), Govt. of India. AM acknowledges CSIR-NET fellowship from the Government of India.

