## Supporting Information for "Connecting Cryo-EM and Crystallographic Views of RNA Folding through Ionic Conditions and Structural Flexibility"

**This PDF file includes:**

**Supporting Information:**

➔ Computational Methods

- All-atom Explicit Solvent Simulation Details
- Structural Topology-Based Electrostatic Model (STEM)
- Folding Free Energy Simulation Method
- Markov State Model
- STEM Simulation Details

➔ Supporting Figures

- Figures S1 to S5

➔ References

**Other Supporting Information for this manuscript includes the following**

Movies S1

### Computational Methods

#### All-atom Explicit Solvent Simulation Details

Our initial Cryo-EM and X-ray crystallography resolved structures of an extensive all-atom level molecular dynamics simulation were carried out both in the presence and absence of $Mg^{2+}$ for 1 μs each, totalling 2 μs of sampling. Simulations were performed implementing AMBER99[1] force field with extensions parmbsc0 and chiOL3[2,3] and with the modified $K^+$ ion[4] parameters to prevent crystallization. Equilibrating the system to acquire the correct distribution of outer-sphere and inner-sphere ions in a combination of ions ($K^+$, $Mg^{2+}$) is a challenging task. At first, the RNA was centred in a cubic box having size 100 Å × 100 Å× 100 Å and was solvated with the TIP3P water model[5]. We added 13 $Mg^{2+}$ ions, 106 $K^+$ ions, and 65 $Cl^-$ions, maintaining the physiological concentration of 2 mM [$Mg^{2+}$] and approximately 100 mM [$K^+$] ions.

We used a combination of excess ions to balance the negative RNA charge and bulk ions to account for the physiological ion concentration range and to prevent them from condensing onto the RNA without a suitable hydration shell. They were initially distributed randomly with greater van der Waals radii[6]. Following that, the entire system was advanced to the energy minimization process using the steepest descent algorithm. Keeping the RNA frozen, the solvent and the ions were then equilibrated for 10 ns using an NVT ensemble in each case. Following our early established protocol, the RNA was gradually released in 4 steps, gradually lowering the position restraining force by 1000, 100, 10, and 0 $kJ/mol/nm^2$ at constant volume, spending 5 ns at each step overall collecting 20 ns of NVT equilibration. Finally, the system was initiated for an unbiased explicit solvent simulation run starting with a short 5 ns NPT equilibration, followed by a 1 μs MD NVT production run. In these simulations, a leapfrog integrator with a time step of 2 fs and a Nose–Hoover[7,8] temperature coupling and the LINCS algorithm[9] to constrain all the hydrogen-atom-mediated covalent bonds were applied. For the NPT equilibration, we used the Parrinello–Rahman barostat[10] to maintain an average pressure of 1 bar, and for creating the long-range electrostatic interactions simulations, the Ewald algorithm was employed with a grid spacing of 1.2 Å and 10 Å as coulomb cutoff and within 10 Å radii of van der Waals cutoff, other nonbonded interactions were taken care of. All the simulations used periodic boundary conditions in all directions.

**Structural Topology-Based Electrostatic Model (STEM)**

The STEM framework involves an implicit-explicit ion environment for monovalent (implicit) and divalent salts (explicit) around RNA. It incorporates the Generalized Manning Counterion Condensation (GMCC) theory[11] into an all-atom Structure-Based Model (SBM) of RNA[12–15]. Because of the implicit treatment of salt buffer, STEM simulations are computationally economical and can efficiently sample a large phase-space of a dynamic system like RNA, along with its dynamic ion-environment.

The STEM addresses physiological conditions where RNA is exposed to a mixed environment of monovalent and divalent salts. Explicit treatment of $Mg^{2+}$ is crucial for managing RNA tertiary packing, whereas KCl is treated implicitly for computational efficiency using GMCC theory. GMCC theory extends the Classical Manning Condensation Theory by considering counterion condensation in polyelectrolyte systems[16,17]. In line with GMCC, the STEM incorporates the local charge density of implicit $K^+$ ions as a smeared Gaussian shell around each negatively charged phosphate group. This Gaussian shell creates the hybrid implicit-explicit interface, excluding the continuum charge density of condensed implicit $K^+$ ions from the excluded volume of the explicit phosphate groups.

In the model, $K^+$ and $Cl^-$ are represented with Gaussian smeared charge, while other ions, including phosphate and $Mg^{2+}$, are treated as point charges for simplicity. The interactions—

point charge-point charge, point charge-Gaussian, and Gaussian-Gaussian—are all described in terms of the Debye-Hückel potential. Three of these interaction terms constitute the overall electrostatic potential,

$$\Phi_{Elec}(r_{ij}, \sigma_k) = \Phi(r_{ij}, 0) + \Phi(r_{ij}, \sigma) + \Phi\left(r_{ij}, \sqrt{{\sigma_1}^2 + {\sigma_2}^2}\right) \quad (1)$$

The details of the STEM have been extensively discussed in our recent work[18,19].

**Folding Free Energy Simulation Method**

To sample the whole conformation landscape of bend and stacked conformation of SARS-CoV-2 RNA FSE, we calculated folding free energy using the umbrella sampling method[20] where the fraction of the global native contacts (Q) is selected as an order parameter. A collection of initial structures is generated for each window of Q by adopting a slow pull criterion, ensuring an ion-equilibrated initial structure. However, we again reinitialized explicit $Mg^{2+}$ ion distribution at each umbrella window to extend the equilibration and ran for 200 million steps. To confirm the substantial overlap in the conformational space, we have used a total of 24 and 34 windows along the Q for XRD and cryo-EM structures, respectively. Next, the weighted histogram analysis method (WHAM)[21] is used to calculate the thermodynamic free energy landscape, G(Q).

**Markov State Model**

Markov state models (MSMs) are widely used in molecular dynamics (MD) studies to identify metastable conformational states, characterize kinetic pathways, and extend accessible timescales by combining information from multiple short trajectories[22,23]. In this work, we constructed MSMs based on simulation trajectories generated using the STEM for both bent and coaxially stacked conformations to resolve their kinetic folding pathways. MSM analysis was carried out using the PyEMMA software package[24].

For MSM construction, the fractions of native contacts associated with stem1, stem2, and stem3 were selected as collective variables. The conformational ensembles sampled in the simulations were then discretized into microstates on the three-dimensional free-energy surface using the K-means clustering algorithm[25]. For both systems studied, the data were partitioned into 100 microstates. A transition probability matrix was subsequently constructed by counting transitions between microstates at a chosen Markovian lag time. Lag-time selection was guided by convergence of implied timescale plots, leading to the choice of a lag time of 60 ps. The resulting microstates were further coarse-grained into kinetically meaningful macrostates using Perron–cluster cluster analysis (PCCA+)[26,27]. Finally, transition path theory (TPT) was employed to identify dominant transition pathways between macrostates[28].

**STEM Simulation Details**

Atomic coordinates and condensation variables ($\mu_{i,s}$ and $\eta_{i,s}$) are evolved with Langevin dynamics with a time step of $0.001\tau_R$. We used an underdamped condition for rapid sampling. For explicit particles, reduced mass of $1\mu_R$ and drag coefficient of 1 ${\tau_R}^{-1}$ are used. Condensation parameters, $\mu_{i,k}$ and $\eta_{i,k}$ are given a mass of 15 $\mu_R$ nm$^2$ and a drag coefficient of 0.05 ${\tau_R}^{-1}$ nm$^2$. Temperature was chosen to $87T_R$ to capture the breathing dynamics. To ready up different $Mg^{2+}$ composition, we created a large cubic box of length 75 nm. The number of $Mg^{2+}$ molecule included in that box determines the overall concentration of the corresponding solutes. Periodic boundary conditions were applied. Each simulation was run with 20 million of time steps. The related parameter set and its calibration are also available in early literature[18,19].

### Figures

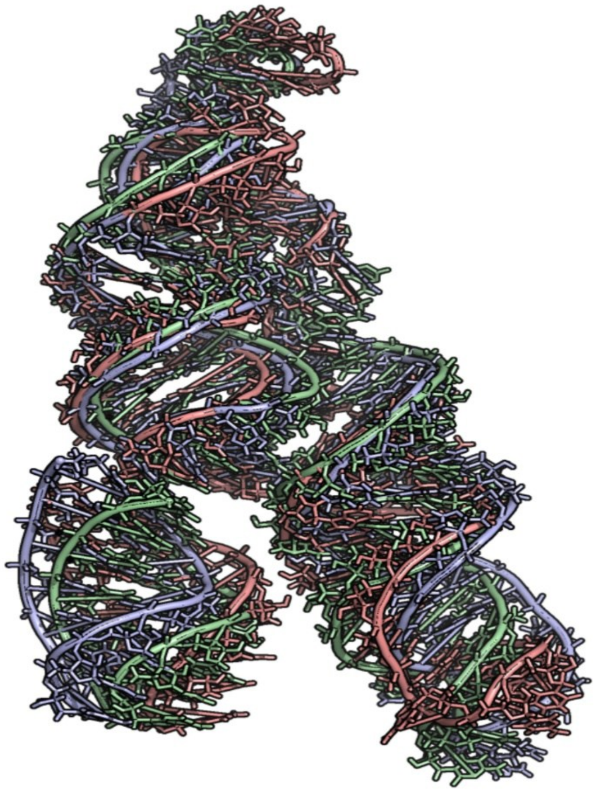

**Figure S1:** Representative conformational ensemble of RNA under no $Mg^{2+}$ conditions.

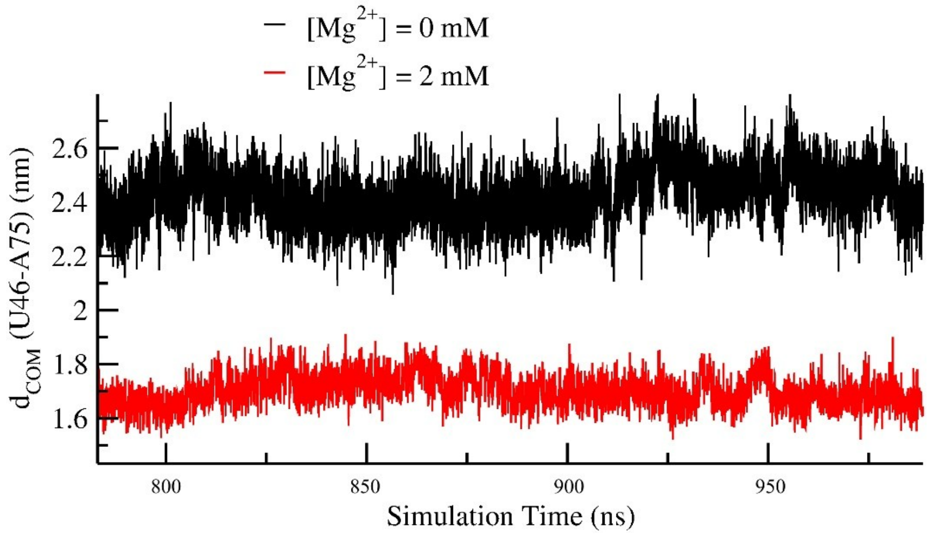


**Figure S2:** $Mg^{2+}$-induced closer approach of the G17–U46–A75 base-triple interaction in the cryo-EM–resolved SARS-CoV-2 FSE. The centre of mass distance between U46 and A75 at $[Mg^{2+}]$ = 0 mM and $[Mg^{2+}]$ = 2 mM conditions.

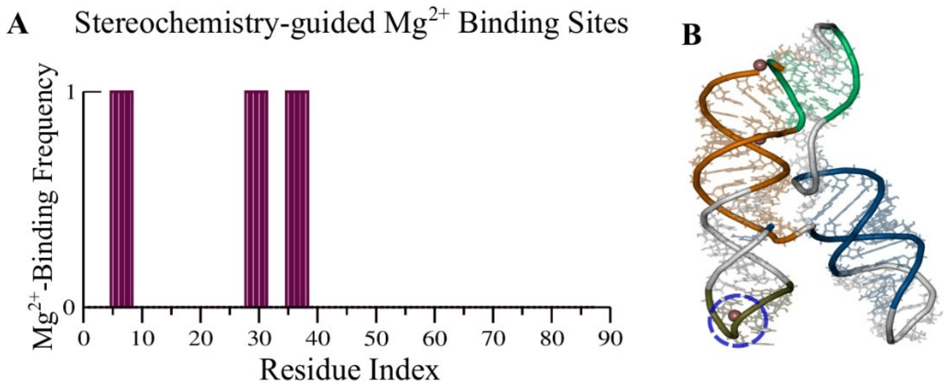


**Figure S3:** Stereochemistry-guided $Mg^{2+}$ binding sites. (A) $Mg^{2+}$ binding sites identified using a stereochemistry-guided method. (B) Representative snapshot of a key $Mg^{2+}$ binding site, with the highlighted region indicating the site driving the conformational transition.

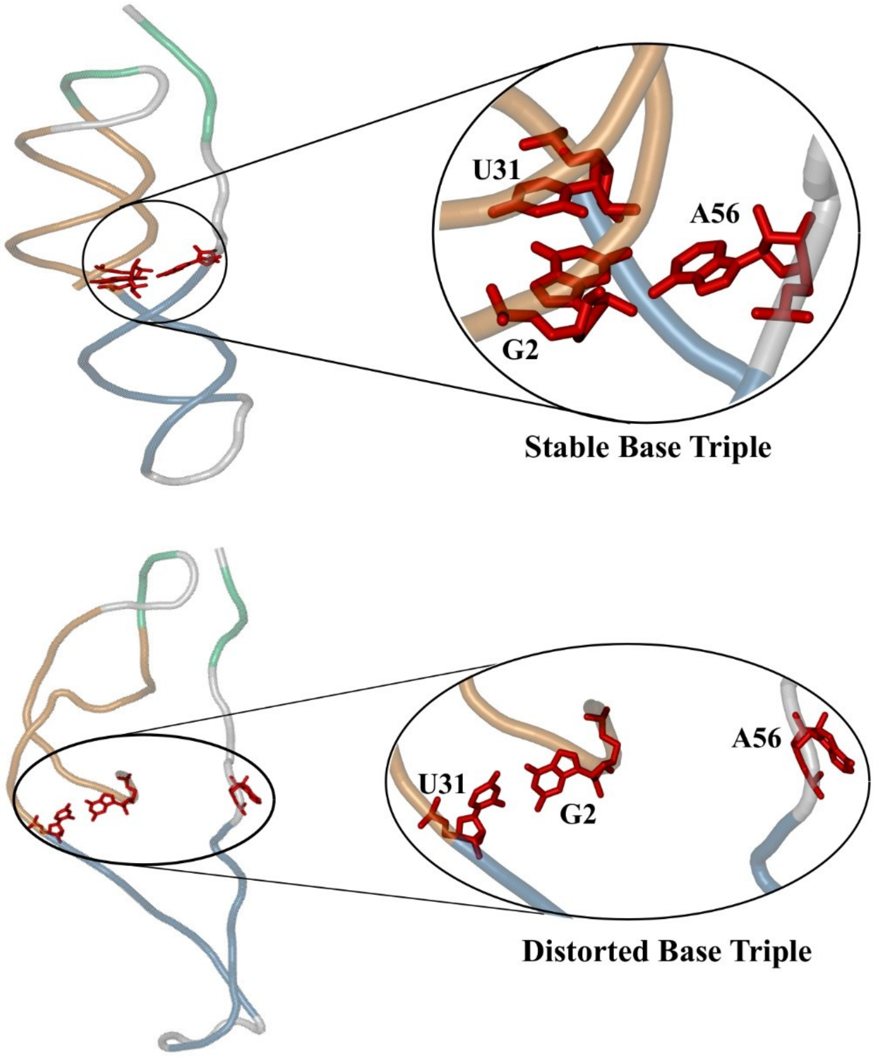


**Figure S4 :** XRD-resolved SARS-CoV-2 RNA captures breathing dynamics that further destabilize the base tripleresponsible for frameshifting.

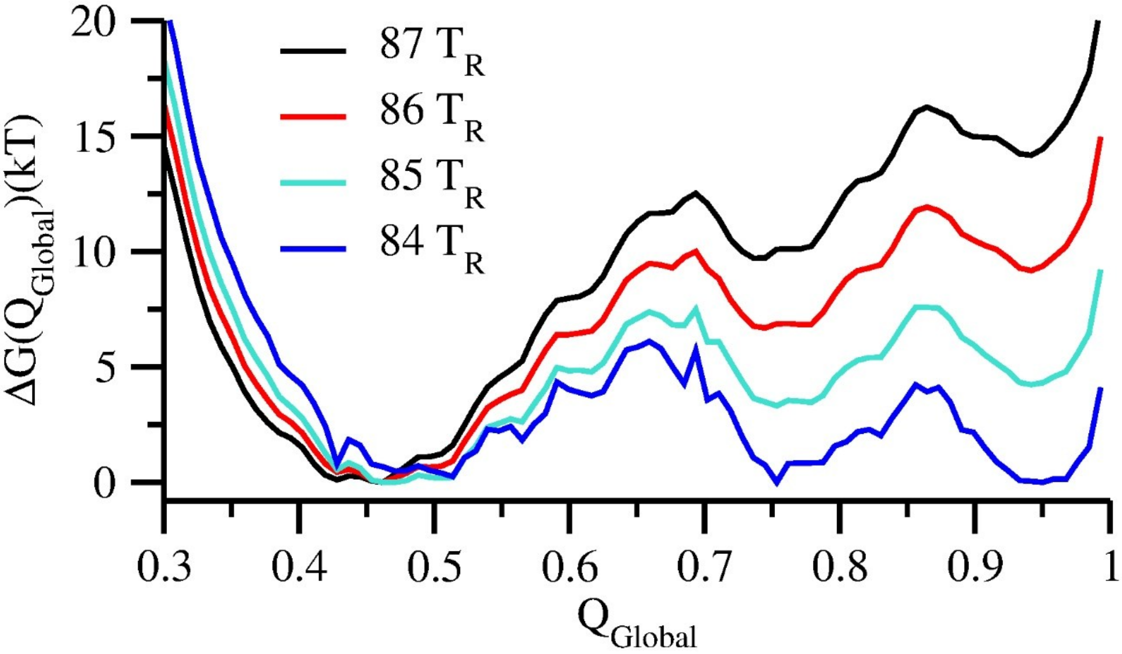


**Figure S5:** Temperature-dependent free energy profiles. Comparison of free energy profiles with different reduced temperatures reveal change in stability between the folding and unfolding states.
